# Accurate estimation of rare cell type fractions from tissue omics data via hierarchical deconvolution

**DOI:** 10.1101/2023.03.15.532820

**Authors:** Penghui Huang, Manqi Cai, Xinghua Lu, Chris McKennan, Jiebiao Wang

## Abstract

Bulk transcriptomics in tissue samples reflects the average expression levels across different cell types and is highly influenced by cellular fractions. As such, it is critical to estimate cellular fractions to both deconfound differential expression analyses and infer cell type-specific differential expression. Since experimentally counting cells is infeasible in most tissues and studies, *in silico* cellular deconvolution methods have been developed as an alternative. However, existing methods are designed for tissues consisting of clearly distinguishable cell types and have difficulties estimating highly correlated or rare cell types. To address this challenge, we propose Hierarchical Deconvolution (HiDecon) that uses single-cell RNA sequencing references and a hierarchical cell type tree, which models the similarities among cell types and cell differentiation relationships, to estimate cellular fractions in bulk data. By coordinating cell fractions across layers of the hierarchical tree, cellular fraction information is passed up and down the tree, which helps correct estimation biases by pooling information across related cell types. The flexible hierarchical tree structure also enables estimating rare cell fractions by splitting the tree to higher resolutions. Through simulations and real data applications with the ground truth of measured cellular fractions, we demonstrate that HiDecon significantly outperforms existing methods and accurately estimates cellular fractions.

## 1 Introduction

Tissue-level gene expression data quantify the average expression across cell types, which is largely affected by the heterogeneity of cell type proportions. The varying cellular fractions across tissue samples can confound tissue-level analyses (Jaffe et al., 2014), potentiate the estimation of cell type-specific differential expression (J. Wang et al., 2021), and can help understand the etiology of disease (Mostafavi et al., 2018). Although biochemical methods like flow cytometry and immunohistochemistry can measure a tissue sample’s cell counts, they are labor-intensive and costly. Thus, *in silico* cellular deconvolution methods have been developed to estimate cellular fractions from bulk tissue data as a cost-effective alternative.

Existing deconvolution methods can be grouped into three categories: unsupervised, semi-supervised, and supervised. Unsupervised deconvolution methods do not require a reference and mimic factor analyses, but the resulting factors are usually hard to annotate and interpret (Avila Cobos et al., 2020). Semi-supervised deconvolution methods depend on marker genes that are only expressed in certain cell types and are used to infer cell type references solely using bulk data (Zhong et al., 2013).With the development of sorted-cell or single-cell RNA sequencing (scRNA-seq) reference data, supervised deconvolution has become a powerful alternative and has precipitated the development of methods that leverage these references to better estimate cell type proportions (Newman et al., 2015; Hunt et al., 2019; X. Wang et al., 2019; Wilson et al., 2020).

As the number of cell clusters obtained from numerous single-cell studies increases, cell type hierarchy has become important for understanding the topology of cell types across datasets (Wu et al., 2020; Peng et al., 2021; L. Chen et al., 2022). Large efforts have been made to enhance the interpretation of cell types, such as cell ontology (Miller et al., 2020) and hierarchically organized cell types (Hodge et al., 2019). In practice, tissues with various differentiated cell types that share the same origin of cell differentiation, like peripheral blood mononuclear cells (PBMCs), bring great difficulties to reference-based deconvolution methods because cell types from the same origin have similar expression levels. This begets co-linearity in cellular deconvolution regression models, which results in highly variable estimates. Some methods attempt to alleviate this issue by selecting better cell type marker genes to reduce the correlation between the reference gene expression levels in different cell types. However, these methods give highly biased results when the quality of cell subtype marker genes is not high (Fischer et al., 2021). In addition, most deconvolution methods work better for more abundant cell types and thus are limited to applications of major cell types. Their estimated fractions of rare cell types are often zero, which precludes downstream analyses of rare but biologically important cell types.

To address the issue of co-linearity and rare types, existing methods HEpiDISH (Zheng et al., 2018) and MuSiC (X. Wang et al., 2019) implemented a top-down recursive deconvolution process guided by a hierarchical cell-type tree. After estimating cell fractions of major cell types in the first layer, they calculate artificial omics of major cell types and use it as the response in reference-based deconvolution to estimate subtype fractions in the second layer, assuming the fraction of a major cell type equals the sum of its subtype fractions. Although this process can be extended to hierarchical trees with more than two layers, the top-down approach may fail when the “parent” cell types are poorly estimated. Importantly, the bias of each layer’s estimation will accumulate and increase the estimation bias of cellular fractions in subsequent layers. Therefore, it is pressing to develop methods that can utilize more complicated tree structures to provide accurate estimates of cellular fractions.

Here we present Hierarchical Deconvolution (HiDecon), a penalized approach with constraints from both “parent” and “children” cell types to make full use of a hierarchical tree structure. The hierarchical tree is readily available from well-studied cell lineages or can be learned from hierarchical clustering of scRNA-seq data (Peng et al., 2021). The tree reflects the similarities between cell types and their differential trajectories. The basic intuition of HiDecon is that there exists a summation relationship between the estimation results of adjacent layers. For instance, after estimating cellular fractions at different resolutions with two deconvolution layers, say lymphocytes (layer 1) and B cells and T cells (layer 2), it will be ideal if the estimated proportion of lymphocytes is the sum of B cell and T cell proportions. If these layers’ estimates do not follow the sum constraints implied by the hierarchical tree, it suggests that estimation bias occurs in certain layers and should be corrected by the estimation results of other layers. To fully use the cell type hierarchy and marker information in different layers, HiDecon implements the sum constraint penalties from the upper and lower layers to aggregate estimates across layers for more accurate cellular fraction estimates. This is especially useful for rare cell types, which may be poorly estimated in other methods that do not use the hierarchical tree.

The remainder of the manuscript is organized as follows. We first introduce our model and estimation algorithm in Section 2. Then in Section 3, we compare HiDecon with existing methods via simulations based on a real scRNA-seq dataset of PBMC from COVID-19 patients and controls. In Section 4, we further benchmark HiDecon in large-scale human blood datasets with measured cell counts. We conclude and summarize our findings in Section 5.

## 2 Methods

### 2.1 A model for deconvolving bulk gene expression

Gene expression levels in bulk tissue samples can be modeled as the weighted sum of cell type-specific expression. A reference-based cellular deconvolution model can be written as

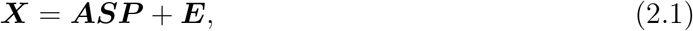

where 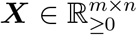 denotes the gene expression levels of *m* marker genes in *n* tissue samples; 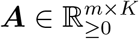 represents the observed reference signature matrix of *K* cell types derived from scRNA-seq or sorted-cell data; 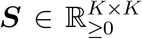 is an observed diagonal matrix with diagonal elements representing the cell size of different cell types (Jia et al., 2017), that is, the average abundance of observed transcripts in each cell type; 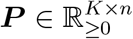 is the cellular fractions for the *K* cell types that need to be estimated; and ***E*** ∈ ℝ^*m*×*n*^ is the error term. Most existing deconvolution algorithms rely on a pre-specified fixed number of cell types (K), but cell types are usually hierarchically organized by cell differentiation or lineage. It is critical to estimate cellular fractions at different resolutions and improve the estimation by borrowing information from related “parent” and “children” cell types. We term the precise cellular deconvolution guided by a hierarchical cell-type tree as hierarchical deconvolution and describe it in detail below.

### 2.2 Hierarchical deconvolution

Assume a known hierarchical tree (either from biology literature or hierarchical clustering) with *L* layers such that cells are split into finer resolutions as *l* ∈ {1,2,…,*L*} increases, where *l* = 0 denotes all cells as a single cluster and *K_l_* is the number of cell types in layer *l*. We first describe how to map cell types across layers *l* and *l* + 1 using a simple example from PBMCs. Assume layer *l* has two clusters representing monocytes and lymphocytes and layer *l* + 1 has three cell types consisting of monocytes, B cells, and T cells, where lymphocytes in layer *l* are divided into B cells and T cells in layer *l* + 1. Let ***p***_*il*_ and ***p***_*i*(*l*+1)_ be sample *i*’s cellular fractions in layers *l* and *l* + 1. Since the fraction of lymphocytes should be similar to the sum of fractions of B cells and T cells, we should have

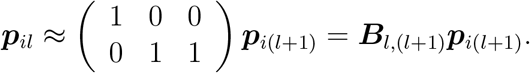

The matrices ***B***_*l*,(*l*+1)_ ∈ {0,1}^*K_l_*×*K*_*l*+1_^ define a mapping between cell types across adjacent layers and parameterize the hierarchical tree, where row k’s non-zero elements are exactly cell type k’s “children” in layer *l* + 1.

We define an estimator for sample i’s cellular fractions from all layers (***p***_*i*1_,…,***p***_*iL*_) to be

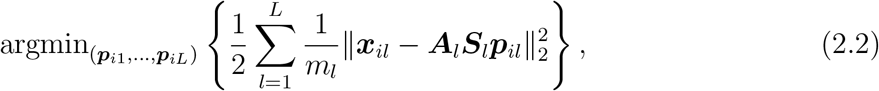

subject to

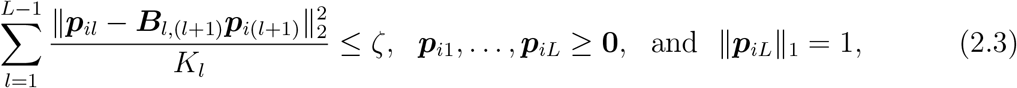

where 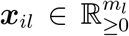 denotes sample *i*’s bulk gene expression at *m_l_* marker genes in layer *l*; 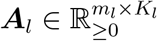 represents the reference signature matrix of *K_l_* cell types derived from scRNA-seq data; and 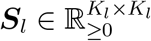 is a diagonal size factor matrix for layer *l* with diagonal entries representing the cell size of the *K_l_* cell types in layer l. Nonnegativity of all parts in the model originates from the nature of gene expression data.

The first constraint in (2.3) reflects the hierarchical tree, and ensures “parent” and “children” cellular fraction estimates are similar. To ensure the interpretation of proportional estimates, we further require the last layer’s fractions to sum to one. The optimization in (2.2) and (2.3) is convex and can be re-written as the following penalized regression problem:

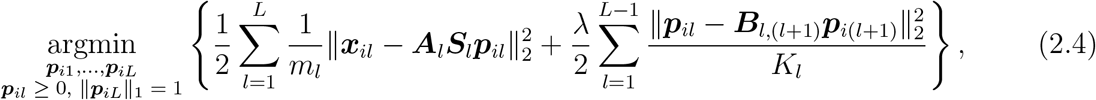

where the tuning parameter λ ≥ 0 is implicitly a decreasing function of *ζ*. That is, λ = 0 implies the tree has no impact on estimates and λ = ∞ means parent cellular fractions are completely determined by their children’s fractions.

### 2.3 Estimation algorithm

To simplify our algorithm and follow the common practice of cellular deconvolution (Mohammadi et al., 2016), we optimize the objective function under the nonnegative constraint and then normalize the fraction estimates for the *L*th layer so they sum to 1. To describe our algorithm, we first note that for 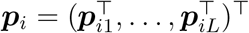, (2.4) can be re-written as the following quadratic problem:

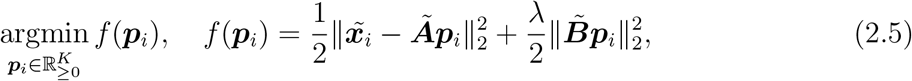

where 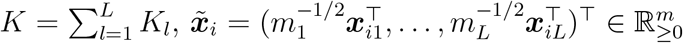, and 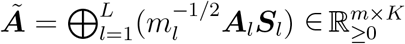 for 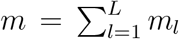. The matrix 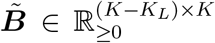 is an upper-triangular difference operator taking the form

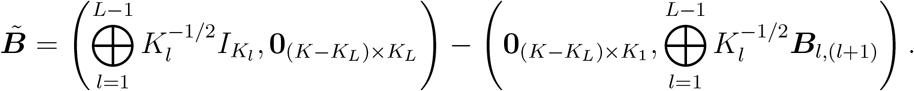

To solve (2.5), we note that the minimizer for ***p***_*i*_’s *k*th coordinate, while fixing all other coordinates, is exactly

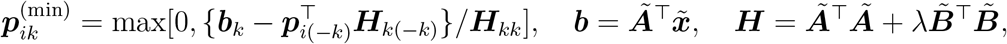

where ***p***_*i*(–*k*)_ and ***H***_(−*k*)*k*_ are the sub-vectors of ***p***_*i*_ and the kth column of ***H*** obtained after deleting their *k*th elements. This naturally leads to Algorithm 1, which employs coordinatewise descent to solve (2.5). While not explicitly stated in Algorithm 1, we normalize our estimate for ***p***_*iL*_, the tree’s last layer’s cellular fractions, so that its entries sum to 1.

**Algorithm 1:**
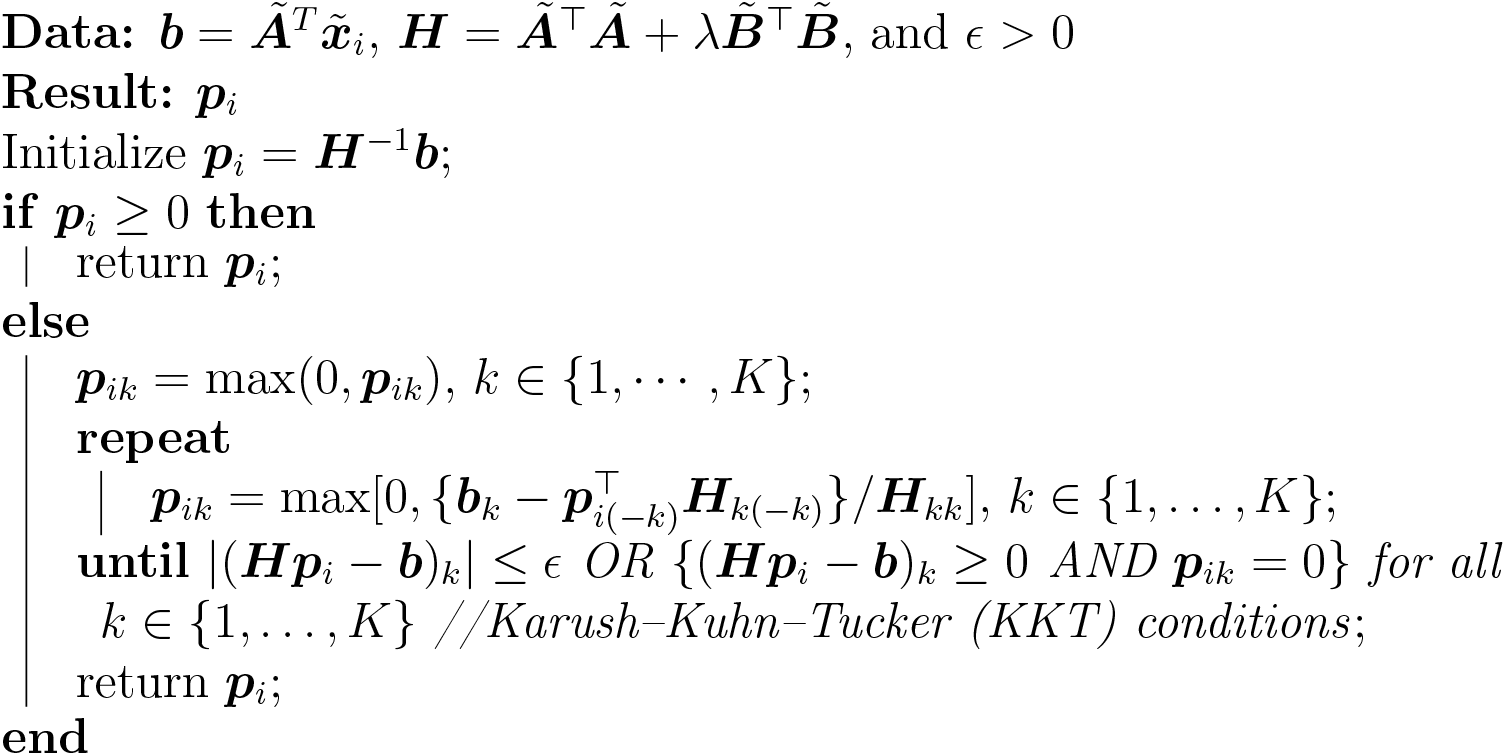
HiDecon optimization algorithm.

### 2.4 Tuning parameter selection

We propose a novel procedure (Figure 1) to select HiDecon’s tuning parameter λ. The idea is to select the optimal λ using a bulk data surrogate with “ground truth” fractions. In order to generate a bulk data surrogate, we apply nonnegative least squares (NNLS) to the observed bulk data 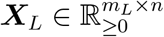 with given signature matrix 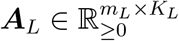,

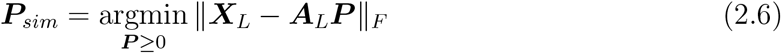

to get rough estimates for cellular fractions ***P***_sim_ of bulk data to imitate the cellular composition structure of tissue samples. Then, we simulate bulk data surrogate ***X***_sim_ with cellular fractions of ***P***_sim_ by resampling individual cells from single-cell reference with replacement. Finally, we compare the performance of HiDecon when deconvolving bulk data surrogate ***X***_sim_ under a series of tuning parameter λ’s. We use Lin’s concordance correlation coefficient (Lin, 1989) between HiDecon estimates ***P***_*HiDecon*_ and the “ground truth” ***P***_sim_ for each cell type as the evaluation metric. The λ with the highest mean concordance correlation coefficient across cell types is considered the optimal tuning parameter for HiDecon.

**Figure 1:**
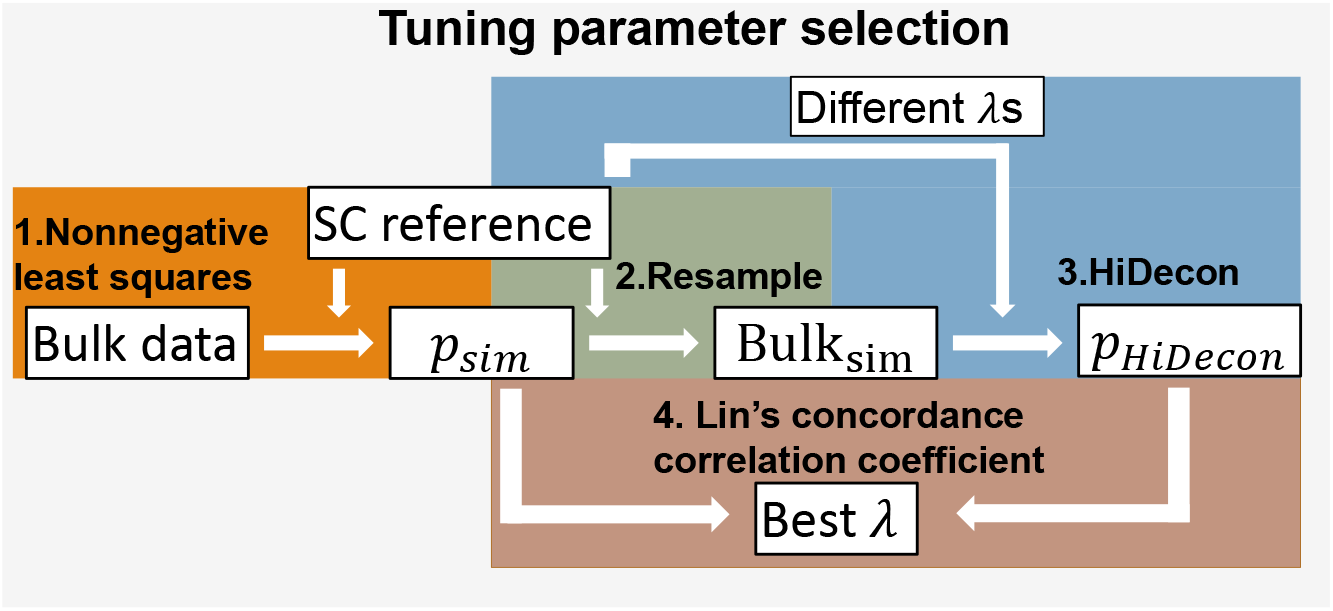
Flow chart for tuning parameter selection. Different steps are marked with different colors.

### 2.5 Data normalization and marker selection methods

We normalize the expression of each tissue sample and cell to the same scale and avoid extreme values. To adjust for library size, the expression of each tissue sample and cell in the raw count matrix is divided by its total count and multiplied by 1,000,000 as the count per million (CPM). Then, data is log_2_ transformed with a pseudo count of 1 to ensure all elements are nonnegative.

In HiDecon, we select marker genes for each layer of the hierarchical tree with the following considerations. First, marker genes selected for coarser clusters are not necessarily marker genes for finer clusters and cannot reduce co-linearity efficiently. Second, in single cell references, coarser clusters might have many cells from subtypes that might not be major in bulk data. Then, the reference mainly represents major cells in the single cell data. Selecting another set of markers for coarser layers can reduce the case that the averaged reference misrepresents the expression level of coarser clusters in bulk data. In both the simulation study and real data application, we use the Wilcoxon rank-sum test to identify the differentially expressed (DE) genes between two cell clusters after normalizing the reference data as described above. To identify DE genes of one cluster compared with all other clusters in this layer, we use the intersection union test (Berger, 1997) to calculate the combined p-value for assigning rankings to genes. We use the top 50 genes with the smallest p-values for each cell cluster as its marker genes.

### 2.6 Evaluation metrics

Denote the ground truth of cell type fractions by ***P***_*K*×*n*_ and the estimated fractions by 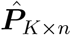. We use two evaluation metrics to evaluate the performances of methods in our simulation study and real data application.

1. Mean absolute error:

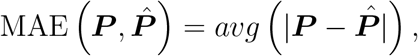

where *avg* of a matrix or a vector is the average over all its entries;
2. Lin’s concordance correlation coefficient (Lin, 1989):

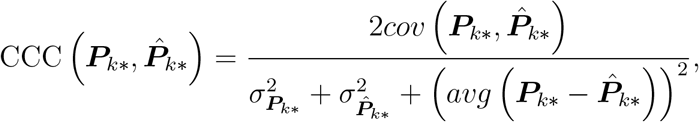

where ***P***_*k*_* denotes the row *k* of **P**, that is, the *k*th cell type, and *σ*^2^ denotes the variance. CCC ∈ (−1,1) is a comprehensive measure sensitive to variability and slope and intercept of the linearity. The concordance improves as the value of CCC approaches 1. It captures the deviation of estimates from ground truth, that is points’ degree of departure from the line *y* = *x*.

## 3 Simulation studies

### 3.1 Simulation benchmarking with large PBMC scRNA-seq data

We first compared HiDecon with existing cellular deconvolution methods via simulations. In addition to the two existing top-down hierarchical deconvolution algorithms (HEpiDISH and MuSiC), we further compared HiDecon with other state-of-the-art deconvolution methods without considering the hierarchical cell tree, including CIBERSORT (Newman et al., 2015) and dtangle (Hunt et al., 2019). We used a real large-scale PBMC scRNA-seq dataset (Ren et al., 2021) to simulate pseudo-bulk data. It is a comprehensive COVID-19 study that contains scRNA-seq data from 27,943 genes of 284 samples, among which there are 28 controls, 122 mild/moderate, and 134 severe/critical samples. When generating bulk data, if there are not enough cells from some cell types, it will introduce large single cell specific variance to the gene expression contribution of this cell type in bulk data. In order to reduce cell specific variance when generating bulk data, we only used samples having at least 20 cells in each type. Moreover, lymphocytes have complicated differentiation structures. We explored subtypes of lymphocyte cell types to evaluate HiDecon’s performance on co-linearity and rare cell types. After filtering samples, we used 608,883 cells from 126 samples to calculate reference and simulate bulk data. Cell types include monocytes (Mono), dendritic cells (DC), B cells (B), natural killer cells (NK), CD4+ T cells (CD4), and CD8+ T cells (CD8). Lymphocyte subtypes (e.g., three subtypes of B cells and CD8+ T cells, respectively) are explicitly shown in Figure 2a, following the cell cluster names in the original paper.

**Figure 2:**
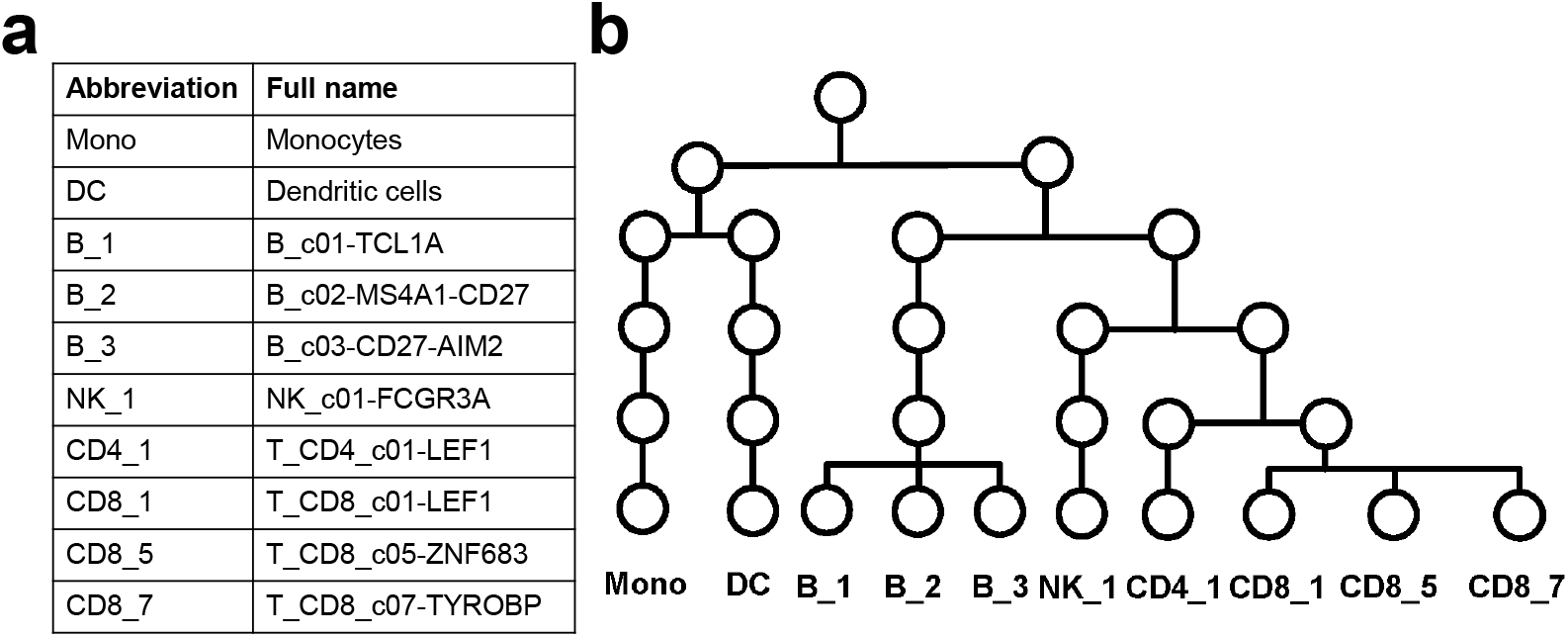
Cell type hierarchical relationship of COVID-19 PBMC data (Ren et al., 2021) used in the simulation. (a) Cell type abbreviation and full name reference. (b) Hierarchical tree constructed from cell lineage relationship and used to guide HiDecon.

We averaged the expression of all cells from each sample to generate simulated bulk data with known fractions, which are calculated by real cell counts of each sample in the scRNA-seq data. It is computationally intensive to select cell type-specific marker genes when the single cell reference matrix has numerous cells. Moreover, having too many cells from certain samples can cover up the information of some other under-represented samples. Considering these problems, first, we averaged single cell gene expression for each cell type to extract sample-level single cell data. This process was performed over all samples. Then, we pooled these sample level single cell data together to construct the pseudo single cell reference so that all samples are equally represented in this reference. The reference dimension is genes by the product of the number of cell types and the number of samples.

We used a hierarchy tree from biological cell lineage relationship (Figure 2b). Dendritic cells are most similar to monocytes. B cells, natural killer cells, CD4 cells, and CD8 cells are all lymphocytes. B cells are mostly naive or memory cells which sets them apart from natural killer cells, CD4 cells, and CD8 cells which are mostly active immune cells. Finally, CD4 cells and CD8 cells are both subtypes of T cells.

We used the tuning parameter selection method to find the best λ for HiDecon. With the ground truth of cellular fractions in the simulated dataset, we calculated CCC between the ground truth fraction ***P***_*K*×*n*_ and deconvolution estimated fraction 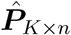 for each cell type across all samples to measure the concordance. We also calculated MAE to assess the accuracy of estimates (Table 1). HEpiDISH can only process the split of one cluster in the first layer, which is not applicable for complex structures and thus not compared in the simulation. HiDecon shows a comprehensively accurate estimation performance with the highest mean CCC and lowest MAE. It is worth noticing that HiDecon is powerful in estimating rare cell type fractions. For the eight cell types with average true abundance lower than 10%, the mean CCC across these types of HiDecon estimates is 0.53 while these of CIBERSORT and dtangle are only 0.36 and 0.42 respectively.

**Table 1:** Comparing the true and estimated cellular fractions in the simulation. The first ten columns show Lin’s concordance correlation coefficient (CCC) calculated for each cell type across all samples. The last two columns present the mean CCC across cell types and MAE (mean absolute errors) for each method. The boldface highlights the best method in each column. Average true fractions for each cell type are shown in parentheses under cell type names. Full names of cell types are listed in Figure 2a.

|  | Mono<br>(0.38) | DC<br>(0.02) | B_1<br>(0.07) | B_2<br>(0.03) | B_3<br>(0.03) | NK_1<br>(0.07) |
| --- | --- | --- | --- | --- | --- | --- |
| HiDecon | 0.85 | 0.25 | **0.80** | **0.54** | **0.50** | 0.34 |
| CIBERSORT | 0.88 | **0.31** | 0.66 | 0.31 | 0.37 | 0.01 |
| dtangle | 0.35 | 0.08 | 0.56 | 0.18 | 0.30 | **0.47** |
| MuSiC | **0.89** | 0.01 | NA | NA | 0.16 | NA |
|  | CD4_1<br>(0.13) | CD8_1<br>(0.09) | CD8_5<br>(0.09) | CD8_7<br>(0.08) | Mean<br>CCC | MAE |
| HiDecon | **0.62** | 0.66 | 0.52 | 0.64 | **0.57** | **0.05** |
| CIBERSORT | 0.42 | **0.76** | 0.25 | 0.25 | 0.42 | 0.07 |
| dtangle | 0.51 | 0.51 | **0.60** | **0.69** | 0.43 | 0.06 |
| MuSiC | NA | 0.35 | 0.22 | 0.39 | 0.20 | 0.08 |

We further checked the estimation details with scatter plots of measured and estimated cellular fractions (Figure 3). HiDecon estimated fractions are well aligned by the diagonal line *y* = *x*, while other methods fail to estimate some cell types. CIBERSORT estimates natural killer cells NK_1 (NK_c01-FCGR3A) as zero in 95.2% of samples. dtangle has generally flat estimates for lymphocyte subtypes. MuSiC has exaggerated estimates for dendritic cells and CD8_1 (T_CD8_c01-LEF1) cells and flat estimates for CD8_5 (T_CD8_c05-ZNF683) cells. In contrast to other methods that have many zero estimates, HiDecon saves many rare cell type fractions from being estimated as zero, providing evidence of HiDecon’s ability to estimate rare cell types.

**Figure 3:**
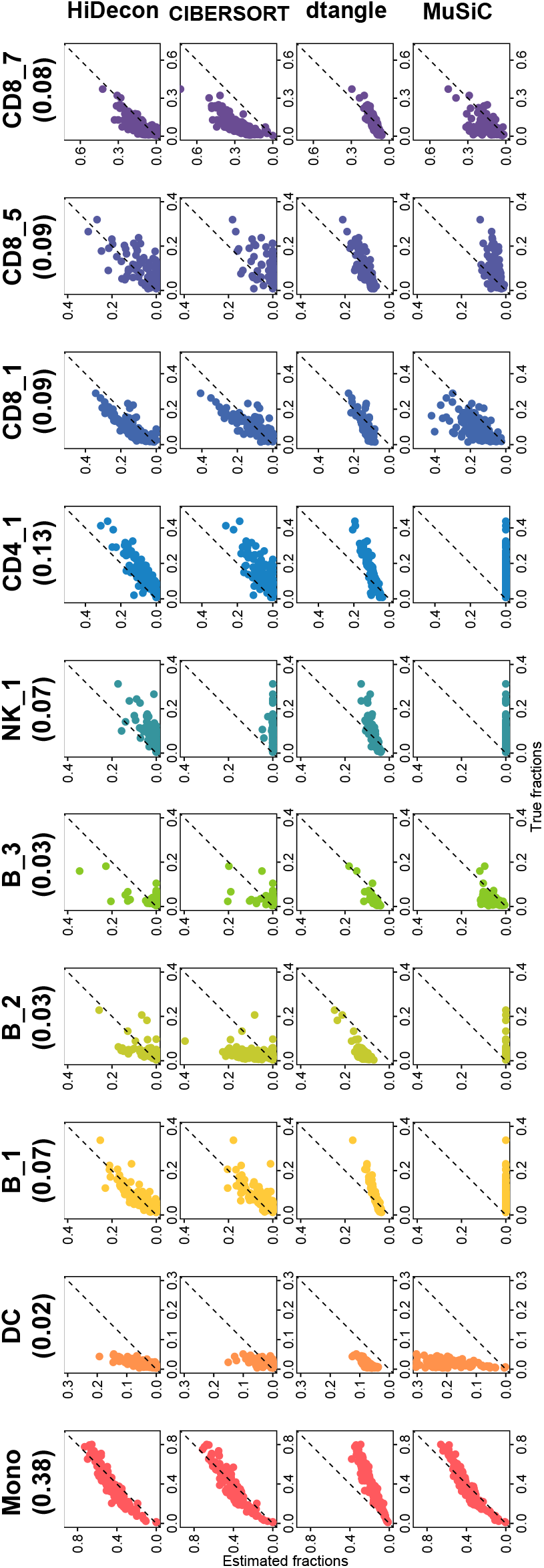
Scatter plots of cellular fractions in the C0VID-19 data simulation study for different deconvolution methods. The x-axis represents the ground truth of PBMC cell fractions, and the y-axis shows the estimated cell type fractions. Cell type names and their mean fractions calculated by ground truth are shown at the top.

### 3.2 Robustness evaluation

In real settings, there is a deviation from weighted signature sum and bulk data, that is, the error term ***E*** shown in Equation 2.1. Here, we added noises to simulated bulk data and kept track of estimation performances as the noise level gets greater. We set noise *e* ~ *N* (0, *sd*^2^), where the error standard deviation (*sd*) ranges from 0 to the standard deviation of the simulated pseudo-bulk data when there is no noise, with a step size of 0.1. For each level of noise, we repeated the experiments 50 times under different random seeds to eliminate the randomness and we averaged evaluation metrics from the 50 repetitions. We calculated the mean concordance correlation coefficient (CCC) and mean absolute error (MAE) trajectories and made box plots under different levels of noise (Figure 4). HiDecon has higher CCC and lower MAE than CIBERSORT, dtangle, and MuSiC. The CCC curve for MuSiC is not shown due to all zero estimates of some cell types, probably caused by the top-down recursive tree-guided process in MuSiC. The experiment shows that HiDecon has consistently outstanding performances under different noise levels.

**Figure 4:**
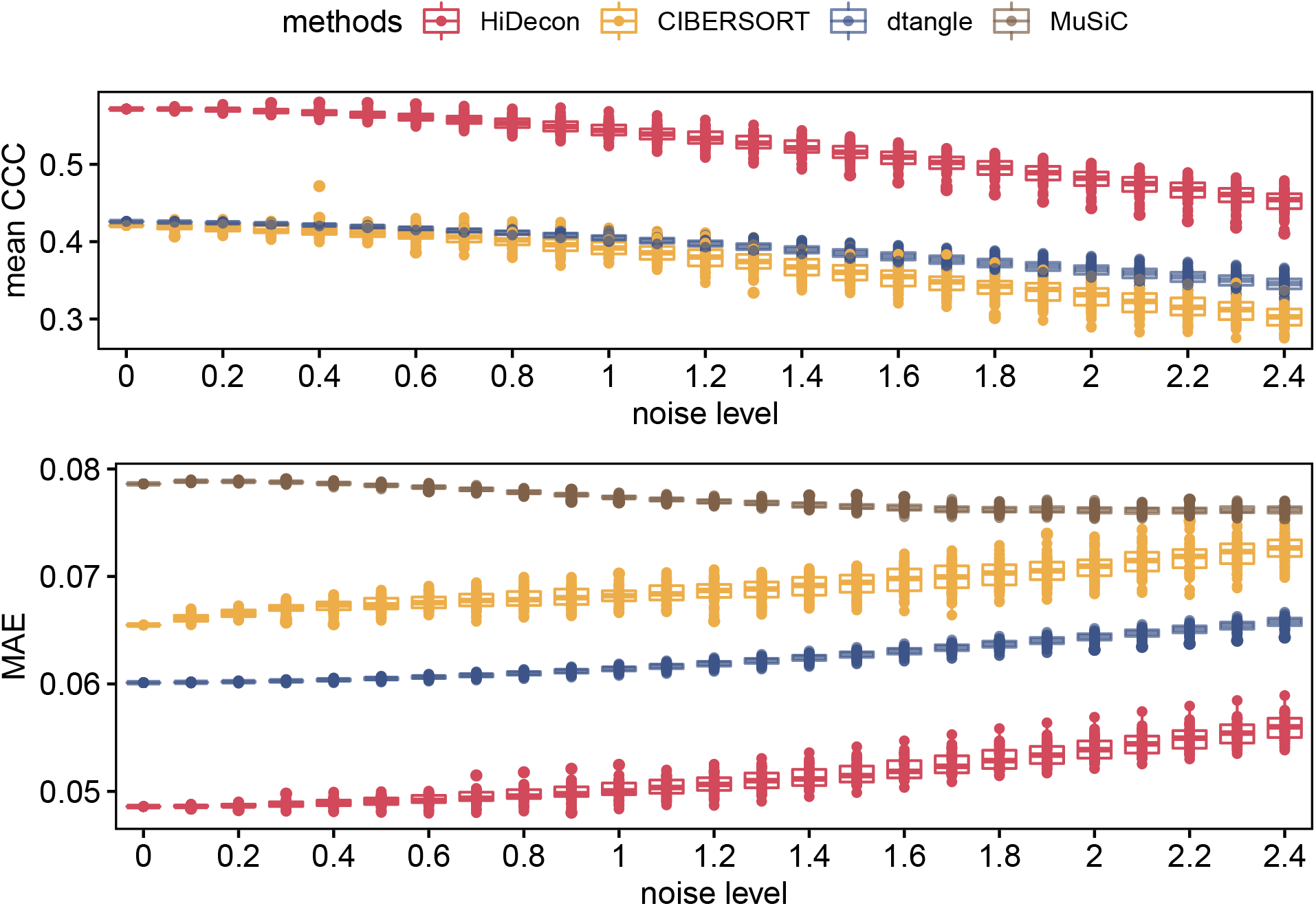
Mean concordance correlation coefficient (CCC) and mean absolute error (MAE) trajectories and box plots under different levels of noises (standard deviation, sd) with 50 repetitions. Noises *e* ~ *N*(0, *sd*^2^) are added to simulated bulk data. The CCC curve for MuSiC is not shown due to all zero estimates of some cell types.

## 4 Real data applications

To assess the performance of HiDecon in real data, we used the Framingham Heart Study (FHS) dataset with measured blood cell counts. FHS is a large-scale longitudinal study with three cohorts: the original (Dawber et al., 1951), offspring (Feinleib et al., 1975), and third-generation cohorts (Splansky et al., 2007). White blood cell (WBC) counts were measured from a complete blood count using the Coulter HmX Hematology Analyzer. There are 4,110 samples from two FHS cohorts (offspring and third-generation) that have both measured cell counts and high-throughput gene expression data from blood. The counted white blood cell types include neutrophils (Neutro), monocytes (Mono), lymphocytes (Lymph), and eosinophils (Eosino). Based on cell lineage, we use a two-layer hierarchical cell tree to guide deconvolution. The second layer has all four cell types, while in the first layer, the lymphocytes are treated as a separate cell type as lymphoid cells, and the other three cell types form a combined cell type to indicate that they belong to myeloid cells.

Similar to the simulation, we used CCC and MAE as evaluation metrics to compare the estimated and measured fractions. As expected, HiDecon demonstrates superior concordance and the lowest MAE across almost all cell types (Table 2). HiDecon’s CCC on eosinophils, a rare type with only 3% abundance, is more than 4 times higher than other methods.

**Table 2:** Lin’s concordance correlation coefficient (CCC) and MAE between measured and estimated cellular fractions in the FHS data. Missing values (NAs) are caused by all zero estimates of some cell types. When calculating the mean CCC, NAs are treated as zero. The boldface highlights the best method in each column. Average true fractions for each cell type are shown in parentheses under cell type names.

|  | Neutrophil<br>(0.60) | Lymphocyte<br>(0.28) | Monocyte<br>(0.09) | Eosinophil<br>(0.03) | Mean<br>CCC | MAE |
| --- | --- | --- | --- | --- | --- | --- |
| HiDecon | 0.13 | **0.57** | **0.04** | **0.28** | **0.26** | **0.10** |
| CIBERSORT | **0.15** | 0.31 | 0.02 | 0.06 | 0.13 | 0.15 |
| dtangle | 0.02 | 0.17 | 0.01 | 0.01 | 0.05 | 0.17 |
| HEpiDISH | 0.12 | 0.25 | 0.03 | 0.04 | 0.11 | 0.17 |
| MuSiC | NA | 0.08 | 0.00 | NA | 0.02 | 0.32 |

As visualized in the scatter plots, HiDecon estimated fractions have a better concordance with measured fractions along the diagonal line than other methods (Figure 5). Methods like CIBERSORT, HEpiDISH, and MuSiC produce many false zeros or even all zero estimates in some cell types, while dtangle shows flat estimated fractions which is similar to that in the simulation scatter plot (Figure 3). Additionally, HEpiDISH and CIBERSORT estimated fractions have exaggerated variability.

**Figure 5:**
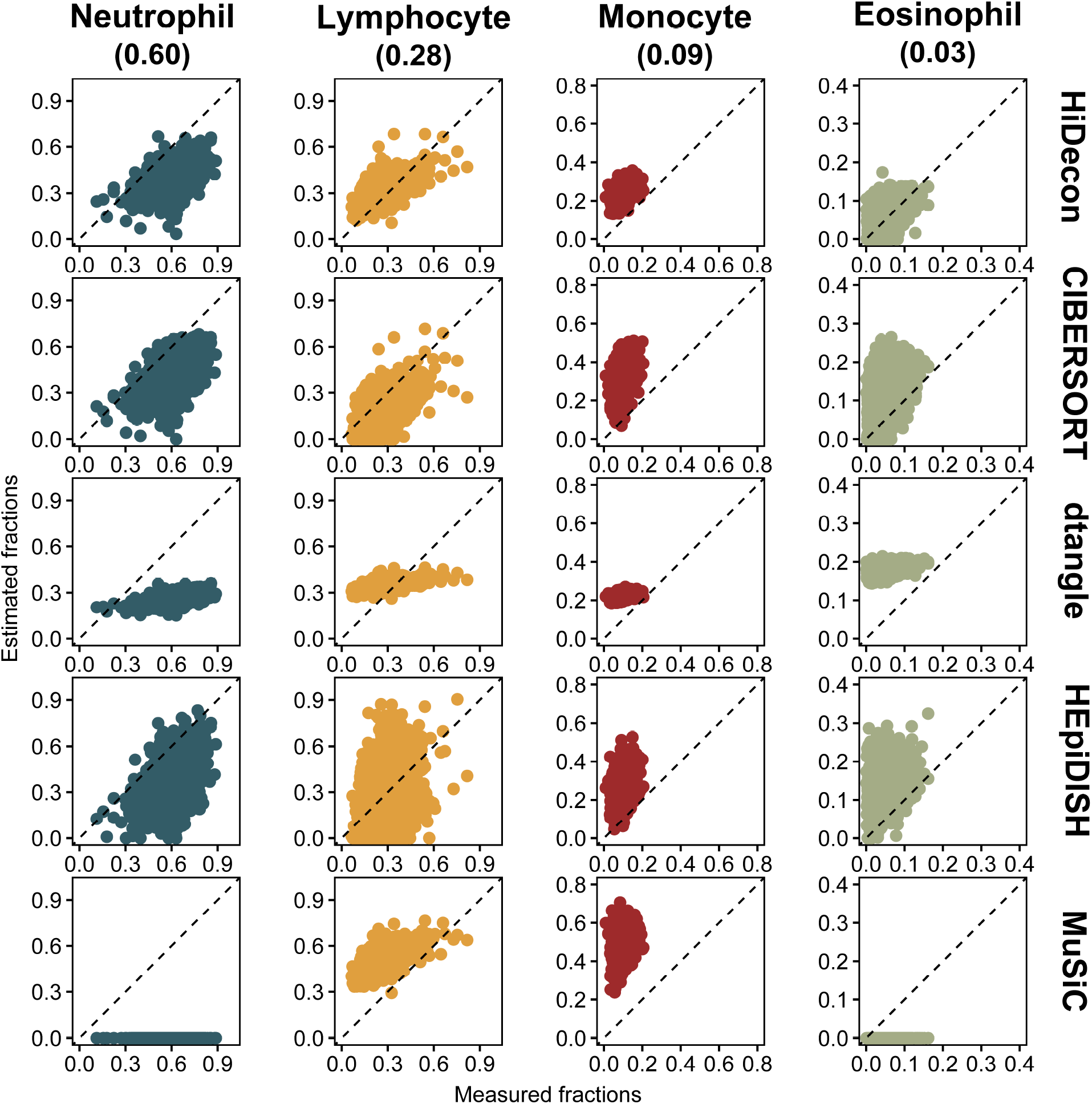
Scatter plots of cellular fractions in the FHS data for different deconvolution methods. The x-axis represents the ground truth of measured white blood cell fractions, and the y-axis shows the estimated cell type fractions. Cell type names and their mean fractions calculated by ground truth are shown at the top.

## 5 Discussion

In summary, we proposed hierarchical deconvolution (HiDecon) to incorporate a hierarchical cell type tree into cellular deconvolution to facilitate the estimation of related and rare cell types. To solve the problem of reference-based deconvolution methods that related cell types cause co-linearity in the signature matrix, we developed an algorithm to leverage the constraints from “parent” and “children” cell types when we iteratively estimate the cellular fractions of all layers of a cell type tree. We benchmarked the HiDecon algorithm on simulated COVID-19 PBMC data and a large real human blood dataset from FHS and evaluated HiDecon by comparing estimation results to the true or measured cell counts. Both numerical and real data benchmarking studies indicate that HiDecon shows higher accuracy than existing methods. We implemented the algorithm as a user-friendly R package HiDecon and hosted it on GitHub (https://github.com/randel/HiDecon). HiDecon can incorporate complex hierarchical cell type tree structure, while the software of the two existing hierarchical deconvolution methods, MuSiC and HEpiDISH, can only incorporate a two-layer tree. Importantly, HiDecon enjoys fast convergence speed brought by the convexity of the objective function. It takes only 12.6 seconds to deconvolve the 4,110 bulk samples of the FHS data.

However, HiDecon also has some limitations. First, in our numerical study, blood data have clearly constructed and biologically and statistically interpretable hierarchical tree. If there does not exist known hierarchical tree for a tissue, we recommend users construct celltype relationships by hierarchical clustering using single-cell data. Second, when deconvolving samples in which cell types are all highly distinguishable, HiDecon might not outperform existing methods because the hierarchical tree cannot further help in this setting. This is rare in practice especially as the research interest goes into refined cell types.

Accurate estimation of cell type proportions can provide novel insights for many downstream analyses at cell type resolution. Representative analyses include differential fraction analysis (M. Cai et al., 2022), cell type-specific (CTS) differential expression (Z. Li et al., 2019; J. Wang et al., 2020; Jin et al., 2021), CTS eQTLs (expression quantitative trait loci) (Westra et al., 2015; J. Wang et al., 2021) when both gene expression and genetic data are available, and CTS gene co-expression networks (S. Chen et al., 2020). Furthermore, hierarchical structures are not limited to gene expression data but also exist in other omics data such as DNA methylation. We will explore more settings where a hierarchical tree can serve as a useful guideline to deconvolve other omics data types that warrant future work.

## Acknowledgement

This research was funded in part through NIH’s R01AG080590, R03OD034501, and R01MH123184, and a grant from the University of Pittsburgh UPMC Health System’s Competitive Medical Research Fund. This research was supported in part by the University of Pittsburgh Center for Research Computing through the resources provided. The Framingham Heart Study is conducted and supported by the National Heart, Lung, and Blood Institute (NHLBI) in collaboration with Boston University (Contract No. N01-HC-25195, HHSN268201500001I and 75N92019D00031). This manuscript was not prepared in collaboration with investigators of the Framingham Heart Study and does not necessarily reflect the opinions or views of the Framingham Heart Study, Boston University, or NHLBI.

